# Within Visit Test-Retest Reliability of EEG Profiles in Children with Autism Spectrum Disorder and Typical Development

**DOI:** 10.1101/834697

**Authors:** April R. Levin, Adam J. Naples, Aaron Wolfe Scheffler, Sara J. Webb, Frederick Shic, Catherine A. Sugar, Michael Murias, Raphael A. Bernier, Katarzyna Chawarska, Geraldine Dawson, Susan Faja, Shafali Jeste, Charles A. Nelson, James C. McPartland, Damla Şentürk, the Autism Biomarkers Consortium for Clinical Trials

## Abstract

Biomarker development is currently a high priority in neurodevelopmental disorder research. For many types of biomarkers (particularly biomarkers of diagnosis), reliability over short time periods is critically important. In the field of autism spectrum disorder (ASD), resting electroencephalography (EEG) power spectral densities (PSD) are well-studied for their potential as biomarkers. Classically, such data have been decomposed into pre-specified frequency bands (e.g., delta, theta, alpha, beta, and gamma). Recent technical advances, such as the Fitting Oscillations and One-Over-F (FOOOF) algorithm, allow for targeted characterization of the features that naturally emerge within an EEG PSD, permitting a more detailed characterization of the frequency band-agnostic shape of each individual’s EEG PSD. Here, using two resting EEGs collected a median of 6 days apart from 22 children with ASD and 25 typically developing (TD) controls during the Feasibility Visit of the Autism Biomarkers Consortium for Clinical Trials, we estimate within visit test-retest reliability based on characterization of the PSD shape in two ways: (1) Using the FOOOF algorithm we estimate six parameters (offset, slope, number of peaks, and amplitude, center frequency and bandwidth of the largest alpha peak) that characterize the shape of the EEG PSD; and (2) using nonparametric functional data analyses, we decompose the shape of the EEG PSD into a reduced set of basis functions that characterize individual power spectrum shapes. We show that individuals exhibit idiosyncratic PSD signatures that are stable over recording sessions using both characterizations. Our data show that EEG activity from a brief two-minute recording provides an efficient window into understanding brain activity at the single-subject level with desirable psychometric characteristics that persist across different analytical decomposition methods. This is a necessary step towards analytical validation of biomarkers based on the EEG PSD, and provides insights into parameters of the PSD that offer short-term reliability (and thus promise as potential biomarkers of trait or diagnosis) versus those that are more variable over the short term (and thus may index state or other rapidly dynamic measures of brain function). Future research should address longer-term stability of the PSD, for purposes such as monitoring development or response to treatment.

## Introduction

Development of translational biomarkers is a crucial step towards clinical trial readiness for neurodevelopmental disorders such as Autism Spectrum Disorder (ASD).^1^ The recent failure of several promising clinical trials^2,3^ underscores the importance of biomarker development, and the need for a range of biomarkers serving a range of purposes. For example, a diagnostic biomarker can confirm presence or absence of a disorder, or identify individuals with a biologically-defined subtype thereof,^4^ in order to guide patient selection for clinical trials. A monitoring biomarker can serially assess the status of a disorder,^4^ and thus measure response to medical therapies or other exposures. The ideal properties of a given biomarker thus depend largely on its context of use. For example, a diagnostic biomarker should not change significantly over a given time window if the biology of the disorder it is indexing has not changed. On the other hand, a monitoring biomarker should change over time in a manner that reflects the biological impact of a medical treatment.

One of the most promising imaging tools for biomarker development in neurodevelopmental disorders is electroencephalography (EEG). EEG is an index of the neural networks that bridge genotype to phenotype across a variety of ages, disorders, and species, and thus offers substantial promise for the development of scalable biomarkers that are relevant to the brain mechanisms underlying ASD.^5,6^ Within EEG, the power spectral density (PSD), which represents the contributions of oscillations at various frequencies to the EEG, offers both diagnostic and monitoring potential. For example, among children with ASD compared to typical development, there is evidence that the resting PSD shows (at a group level) excessive power in the low (delta, theta) and high (beta, gamma) frequency bands and insufficient power in the middle (alpha) frequency bands^7^ This suggests potential utility of some aspects of the PSD as a diagnostic biomarker for autism. Moreover, EEG is a measure of cortical activity and is thus fundamentally dynamic; it changes throughout development, across awake and asleep states, and in response to pharmacological treatment. This suggests that there may be aspects of the PSD that offer potential in other categories of biomarker development (e.g., monitoring or response biomarkers).

Thus, to inform the development of biomarkers using EEG-based measures, it is necessary to evaluate the reliability of the PSD within an individual over brief time intervals, as well as across development and in response to various therapies. This is of particular importance in ASD, given the suggestion that intra-individual variability in brain activity may itself be an endophenotype of ASD.^8^ Different features of the PSD may exhibit different measurement properties, with some parameters reflecting more transient or “state-like” properties of brain activity and others reflecting more stable “trait-like” interindividual differences. To begin this process, in the present study, we focus on test-retest reliability of the PSD and specific parameters thereof over a short time window (median of 6 days) during which one would not expect significant changes in underlying diagnosis, developmental changes are minimal, no new treatments are given, and EEG is collected under identical conditions.

Prior studies in healthy adults have demonstrated good to excellent test-retest reliability for certain features of the PSD. EEG power for mid-range frequencies (theta, alpha, and beta, as opposed to delta and gamma)^9^ and relative power (as opposed to absolute power)^10^ have shown correlation coefficients >.8 for EEG sessions a few weeks apart; this is in the range of test-retest correlations for commonly used tests of cognitive ability.^11,12^ Methodological advances in EEG pre-processing, such as robust reference to average and wavelet independent component analysis which act to attenuate the effects of data collection artifact, improve test-retest reliability in higher frequency bands such as beta and gamma.^13^ However, the reliability of these features in children with or without neurodevelopmental disabilities remains unmeasured.

Notably, traditional methods of characterizing the PSD rely on measuring power within a particular frequency band, which conflates important aspects of underlying EEG activity. First, the EEG PSD typically contains a series of periodic oscillations atop an aperiodic background activity in which the power decreases as frequency (f) increases, leading to a consistent 1/f^α^ distribution to the PSD, with the exponent α determining the slope of this background activity. This aperiodic activity, and the offset thereof, may reflect crucial mechanistic underpinnings of brain activity,^14^ such as tonic excitation/inhibition balance or total spiking activity of underlying neural populations respectively.^15^ The influence of this background activity on the measurement of oscillatory activity is partially (though not completely) eliminated using techniques such as normalization or log transform of the PSD. Second, a priori assumptions about the frequency bands wherein oscillations occur may actually compromise accurate measurement and fail to capture meaningful variation of these oscillations. For example, averaging power in the predefined alpha range (e.g., 8-13 Hz) removes information about the peak alpha frequency in a given individual; however, the exact location of this alpha peak is well known to change with age and cognitive status^16,17^ and can even occur outside of the 8-13 Hz range. Because oscillations rarely span the exact range specified in a frequency band, their activity can be inadvertently included in neighboring frequency bands if they are wide or shifted. Finally, in cases where a periodic oscillation has a narrow bandwidth or is nonexistent a prespecified frequency band, measurement of activity in that band will predominantly reflect aperiodic activity. For these reasons, it is useful to characterize the EEG as a unique profile, with parameterization informed by the shape of each individual’s PSD rather than piecemeal averages across distinct frequency bands.

As of October 2019 ClinicalTrials.gov reported 315 currently recruiting studies collecting EEG data and of those 102 were recruiting pediatric populations. Given the extent of this ongoing research, addressing how best to characterize the profile of the EEG PSD and determine its reliability and stability over time, particularly in clinical and developmental populations, is both important and timely. Such work forms an important foundation on which to base future research, and provides critical information to contextualize current findings.

In this study we therefore explore the test-retest reliability of the profile of the EEG PSD in children with ASD and typical development (TD) over EEG recordings conducted within a short (∼6 day) time-span. We applied two approaches to characterizing the profile of the PSD: (1) parametric model-based decomposition of the PSD into offset, slope, and oscillatory peaks using the Fitting Oscillations and One-Over-F (FOOOF) algorithm^15^; and (2) nonparametric functional data analysis, which identifies a small set of principal component functions that combine to describe the shape of the We hypothesized that these complementary approaches would exhibit high levels of short-term test-retest reliability. In this way, we demonstrate the utility of resting EEG PSD shape, and some specific parameters thereof, as stable biomarkers of cortical activity over short time windows.

## Materials and Methods

These data were collected as part of the ongoing Autism Biomarkers Consortium for Clinical Trials (ABC-CT; www.asdbiomarkers.org).^18^ The objective of the ABC-CT is to evaluate a set of electrophysiological (EEG), eye-tracking, and behavioral measures for use in clinical trials for ASD. The ABC-CT began with a “Feasibility Study Visit,” which included the participants described below and involved two EEGs separated by a short window of time (median 6 days) as described below. The ABC-CT then moved on to the “Main Study Visits,” which included a larger number of participants, with EEGs separated by longer windows of time (6 weeks, and then 6 months). Only the data from the “Feasibility Study” is included here, as the focus of this manuscript is on the shorter-term test-retest reliability of the EEG PSD; this type of information (two EEGs separated by a few days) was not collected in the “Main Study.” This study was carried out in accordance with the recommendations of the central Institutional Review Board at Yale University, with written informed consent from a parent or legal guardian and assent from each child prior to their participation in the study.

### Participants

51 participants (25 with ASD, 26 with TD), aged 4 to 11 years, were enrolled in the feasibility phase of the ABC-CT; group characteristics are presented in Table 1. Groups differed significantly on age (t(45) = 2.3, *p =* .025) and IQ (t(45) = 4.6, *p* <.001) The “Feasibility Study Visit” consisted of two EEGs on two separate days (termed here “Day 1” and “Day 2”), separated by a short window of time (range 1-22 days, median 6 days) during this phase. Participants were characterized using rigorous autism diagnostic standardized measures (Autism Diagnostic Observation Schedule, 2^nd^ edition (ADOS-2),^19^ Autism Diagnostic Interview - Revised (ADI-R),^20^ and Diagnostic and Statistical Manual of Mental Disorders (DSM-5) criteria^21^) by research-reliable clinicians^22^, and cognitive measures Differential Ability Scales 2^nd^ edition (DAS-II).^23^

**Table 1:** Participant sex, age, and IQ by diagnostic group. * indicates measures that differ by group, as described in the text.

| GROUP | N (N FEMALE) | MEAN AGE (Y) | MIN. AGE (Y) | MAX. AGE (Y) | MEAN IQ (SD) |
| --- | --- | --- | --- | --- | --- |
| ASD | 24 (5) | 8.0* | 4.42 | 11.4 | 95 (21.2)* |
| TD | 26 (9) | 6.6 | 4.01 | 11.4 | 120 (12.4) |

### EEG Protocol

In the feasibility phase of the ABC-CT, EEG acquisition included 6 paradigms,^24^ with “Resting EEG eyes open during calm viewing” of silent, chromatic digital videos (similar to screensavers) collected twice on two separate days. Video stimuli consisted of six 30 second non-social abstract videos purchased from Shutterstock, which were presented to the participant in random order in 3 blocks of 1 minute on each day.^25^ The videos were played forward for 15 seconds and then reversed for the following 15 seconds. To allow for counterbalancing of the methods used in the ABC-CT (Eye Tracking and EEG), at screening, participants were stratified based on variables that could be assessed by phone to include group (ASD/TD), biological sex (male/female), age (split at 8 years 6 months), and cognitive ability (ASD only, assessed in person by a trained clinician at first visit). Half of the participants received eye tracking first at each visit and the other half received EEG first.

All sites had a high density EEG acquisition system (Philips Neuro, Eugene, OR), including either Net Amps 300 (Boston Children’s Hospital, University of California Los Angeles, University of Washington, and Yale University) or Net Amps 400 amplifiers (Duke University). All sites used the 128 electrode HydroCel Geodesic Sensor Nets, applied according to Philips Neuro/Electrical Geodesics, Inc. standards. Four of the five sites removed electrodes 125-128, which are positioned on the participant’s face, from the EEG caps to tolerability of wearing the cap. Appropriate EEG acquisition protocols and software (500Hz sampling rate, MFF file format, onset recording of amplifier and impedance calibrations) were provided to each site. EPrime 2.0 (Psychological Software Tools, Sharpsburg, PA) was used for experimental control. The coordinating site reviewed and provided feedback on net application, adherence to administration protocol, and data quality for every session. Sites conducted regular monthly checks of equipment function.

One participant with ASD refused to wear the net; EEG data was therefore available on 24 ASD and 26 TD participants. After the preprocessing described below, EEG from one additional ASD participant was excluded from the parametric and nonparametric data analyses due to having a substantially lower number of observed segments than the rest of the sample (61 segments versus an average of 91 segments) and only one day of EEG recording. Thus, in total, there was usable data on at least one day from 23 ASD and 26 TD participants. Data on and additional one ASD and one TD participant were recorded only on day 1. There was thus usable data on both days from 22 ASD and 25 TD participants.

### Pre Processing of the EEG

Processing of the raw EEG data was done using the Harvard Automated Processing Pipeline for Electroencephalography (HAPPE)^26^ embedded within the Batch EEG Automated Processing Platform (BEAPP).^27^ In brief, data were 1 Hz high pass and 100 Hz low pass filtered, down sampled to 250 Hz, and run through the HAPPE module including selection of 18 channels corresponding to the 10-20 system channels (excluding Cz, as data were originally collected in reference to Cz), 60 Hz electrical line noise removal, bad channel rejection, wavelet-enhanced thresholding, independent component analysis with automated component rejection,^28,29^ automated segment rejection, interpolation of bad channels, and re-referencing to average. Data were then segmented into two second segments, and the PSD was calculated via multitaper spectral analysis^30,31^ using three tapers. The PSD was estimated for each participant and electrode by averaging the PSDs of artifact free segments. Scalp-wide spectral densities were obtained by averaging spectral densities across the 18 electrodes for each subject on each day.

### Parametric Decomposition of Periodic and Aperiodic Activity

In order to characterize periodic and aperiodic features of the PSD profile, we used the Fitting Oscillations and One-Over-F (FOOOF) algorithm.^15^ The algorithm operates by removing an aperiodic slope (Figure 1) from the absolute PSD in the semilog-power space (linear frequencies and logged power), which is fully characterized by offset and slope terms. After removing the aperiodic component, the spectral density contains periodic oscillatory peaks that are modeled as a finite sum of Gaussians. Each Gaussian peak is defined by its amplitude, center frequency, and bandwidth. Thus, the PSD profile, including both the aperiodic background and periodic oscillations, can be fully parameterized by the following parameters: offset, slope, number of peaks (Gaussians), and the center frequency, amplitude, and bandwidth for each peak. These scalar features are then available for analysis across recording sessions using standard statistical techniques. The FOOOF model parameters were chosen by visually inspecting model fit across a range of parameters, blind to participant group and recording session, and selecting those which best captured oscillatory peaks across all of the recordings. A single parameter set was selected for all recordings, Specifically, the peak bandwidth of oscillatory peaks ranged between 1 and 10 Hz, and the minimum peak height (to be included in the fit) was 1.85 standard deviations above the aperiodic background activity.

**Figure 1:**
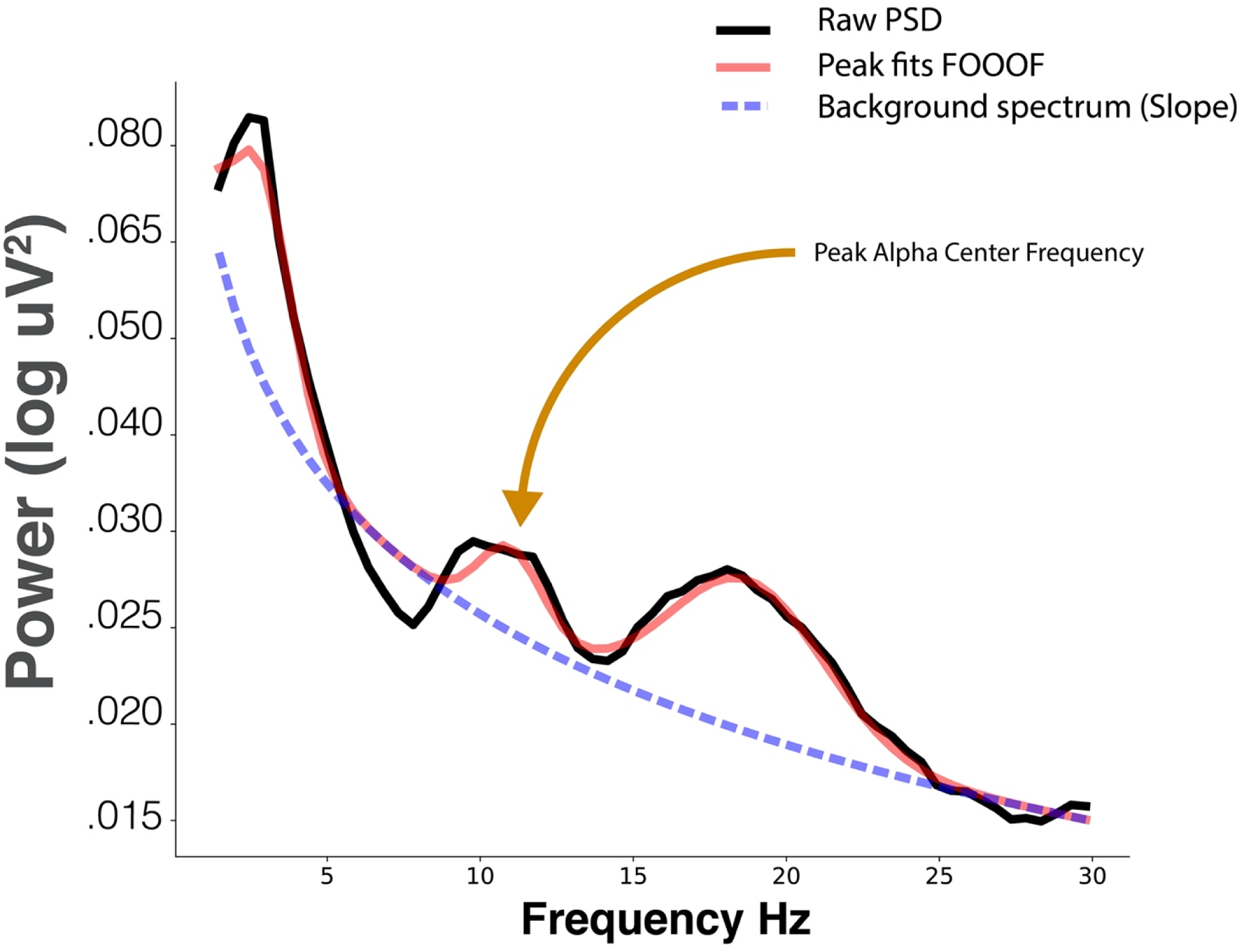
Parameters extracted from FOOOF decomposition of the PSD. FOOOF models individual oscillatory peaks atop the PSD and estimates the slope and offset of aperiodic activity below those peaks.

Since the number of total peaks identified on each spectral density varied across subjects and days, for comparison purposes across consecutive days we first considered the agreement of the location (in terms of frequency band, i.e. delta [2-4 Hz], theta [4-6 Hz], low alpha [6-9 Hz], high alpha [9-13 Hz], beta [13-30 Hz], and gamma [30-55 Hz]) of the peak with the largest amplitude between days. For comparison of the largest peak features (center frequency, amplitude, and bandwidth), we then considered the largest peak in the entire alpha band for stability of results and ease of comparison between diagnostic groups. This allowed characterization of each scalp-wide spectral density by six FOOOF parameters: offset, slope, number of peaks, and (for the largest peak in the alpha range) center frequency, amplitude, and bandwidth. The agreement of these six FOOOF parameters across the two days for each diagnostic group was evaluated using the intraclass correlation coefficient (the ratio of between person variance to total variance) (ICC).^32^ ICC values less than .40 are considered poor, between .40 and .59 fair, between .60 and .74 good, and between .75 and 1.00 excellent.^33^ For all reported ICC values, bootstrap based on resampling subjects with replacement was used for forming percentile confidence intervals (CI). Bootstrap methods yield more reliable inference in small samples (bootstrap CIs were based on 200 resampled data sets).

### Nonparametric Analysis of the Relative Spectral Density via Functional Data Analysis

Scalp-wide relative spectral densities were obtained by averaging relative spectral densities across electrodes for each subject observed on each day. The agreement in relative spectral density across days for both electrode-specific and scalp-wide relative spectral densities was computed by functional ICC within each diagnostic group. Since a trend of lower functional ICC was observed for the most peripheral electrodes (electrodes 9 [FP2], 22 [FP1], 45 [T3], 70 [O1], 83 [O2] and 108 [T4]) across diagnostic groups, a sensitivity analysis was also run through the functional ICC of the scalp-wide relative spectral densities excluding these six electrodes. Computation of functional ICC follows a functional ANOVA decomposition of the data within each diagnostic group. Days are the within subject factor, where the functional ICC can be interpreted as the inter-subject correlation of the entire relative spectral density across days. The functional ANOVA model is fit using a multilevel functional principal components decomposition^34^ which entails estimation of subject- and day-level eigenvalues and eigenfunctions that enrich interpretations by allowing us to connect the nonparametric functional data analysis to results from the parametric analysis via FOOOF. For all reported functional ICC values, bootstrap percentile CIs were formed based on 200 resampled data sets based on resampling from subjects with replacement.

## Results

Age, sex, and IQ for study participants is in Table 1.

The power spectrum of each individual on day 1 and day 2 is plotted in Figure 2. Within participant PSD shapes exhibit striking visual similarity across separate recording sessions.

**Figure 2:**
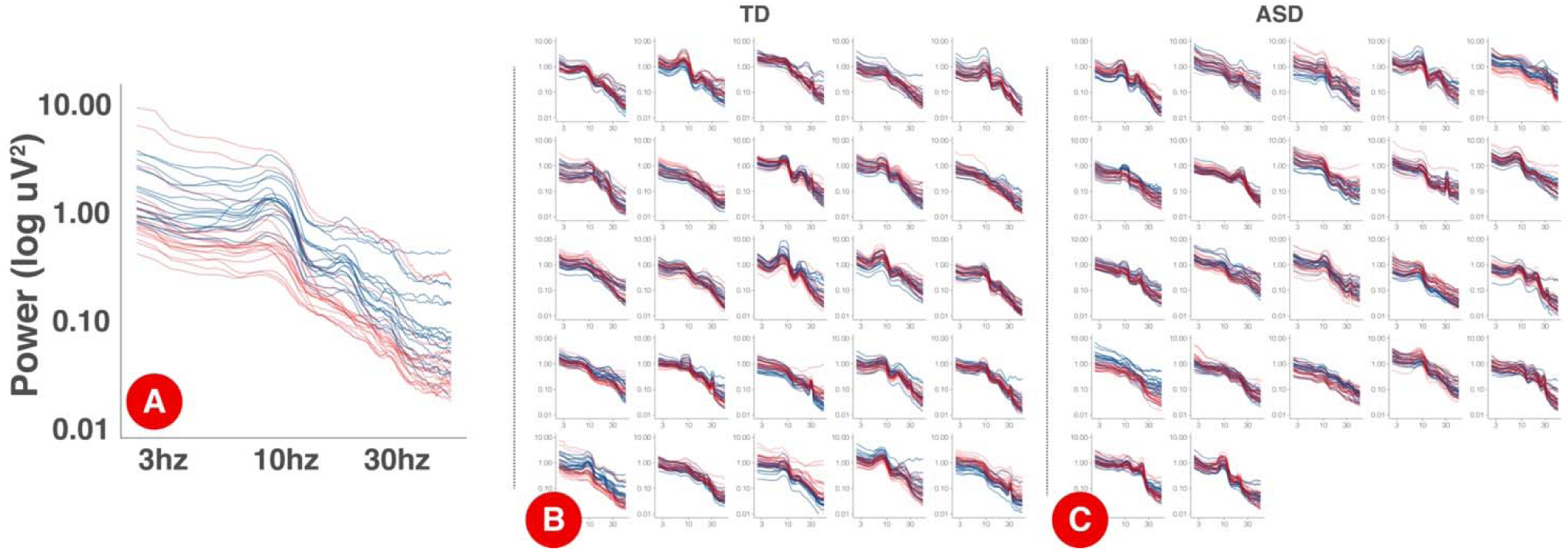
PSDs for each session by participant. Panel A displays an expanded, single participant, PSD with the log-10 axis labels. Each electrode is a single line. Day one PSDs are shown in blue and day 2 PSDs are shown in red. Panels B and C show individual PSDs for TD (B) and ASD (C) participants. Each smaller figure is data from a single participant.

Data quality metrics output from HAPPE^26^ are described in Table 2. Overall, data quality was high across groups.

**Table 2:** Data quality measures, based on HAPPE metrics. Data are reported as mean (SD). EEG segments are 2 seconds long.

| Group | Day | Good Channels (%) | # of EEG segments retained | Rejected components (%) | EEG variance retained (%) | Mean retained artifact probability | Median retained artifact probability |
| --- | --- | --- | --- | --- | --- | --- | --- |
| ASD | 1 | 95.4 (3.4) | 90.7 (1.8) | 29 (11) | 70.2 (17.1) | 0.08 (0.03) | 0.03 (0.02) |
|  | 2 | 95.9 (3.9) | 90.7 (1.8) | 30 (12) | 70.6 (15.8) | 0.08 (0.03) | 0.02 (0.02) |
| TD | 1 | 97.4 (3.8) | 90.8 (1.7) | 18 (10) | 82.5 (13.2) | 0.05 (0.02) | 0.01 (0.01) |
|  | 2 | 97.1 (3.8) | 90.9 (1.7) | 19 (10) | 80.2 (15.2) | 0.06 (0.04) | 0.02 (0.02) |

### Parametric Analysis of the Absolute Power Spectral Density via FOOOF

The location of the dominant peak (i.e. the peak with the greatest amplitude according to the FOOOF algorithm) from both days are provided in Table 3 for both diagnostic groups. The dominant peak occurred most frequently in the high alpha frequency band in the ASD group and low alpha frequency band in the TD group. Across days, while the dominant peak stayed within the alpha band (low and high alpha) mostly for the TD group, it stayed more broadly within the alpha-beta range in the ASD group.

**Table 3:** The location of the dominant peak in day 1 (rows) versus day 2 (columns) among the TD and ASD groups. Values indicate the number of participants with a given combination of dominant peak locations across days.

| TD |  |  |  |  |
| --- | --- | --- | --- | --- |
| Day 1/2 | low_alpha | high_alpha | beta | gamma |
| low_alpha | 6 | 6 | 0 | 0 |
| high_alpha | 5 | 3 | 0 | 1 |
| beta | 1 | 1 | 0 | 0 |
| gamma | 1 | 0 | 0 | 1 |

| ASD |  |  |  |  |
| --- | --- | --- | --- | --- |
| Day 1/2 | low_alpha | high_alpha | beta | gamma |
| low_alpha | 2 | 2 | 1 | 0 |
| high_alpha | 2 | 4 | 3 | 0 |
| beta | 2 | 3 | 1 | 0 |
| gamma | 0 | 1 | 1 | 0 |

The estimated ICCs along with their bootstrap CIs for agreement of the six FOOOF parameters derived from scalp-wide absolute PSD across the two experimental days are provided in Table 4 for both diagnostic groups. Among offset, slope, and number of peaks, offset yielded consistently fair agreement in both groups (TD 0.484 95% CI [0.004, 0.775]; ASD 0.525 95% CI [0.167, 0.806]), with slope between the two days showing poor agreement in the TD group (0.284 95% CI [0,0.674] but good agreement in the ASD group (0.699 95% CI [0.527, 0.815]). Among the three FOOOF parameters describing the largest alpha peak, amplitude had the highest ICC in both groups (TD 0.862 95% CI [0.729, 0.939]; ASD 0.828 95% CI [0.664, 0.926]), followed by center frequency (TD 0.700 95% CI [0.437, 0.862]; ASD 0.619 95% CI [0.342, 0.852]), and bandwidth (TD 0.424 95% CI [0.028, 0.696]; ASD 0.340 95% CI [0.034, 0.727]). While the agreement of the largest alpha peak amplitude was high in both groups, agreement in the peak frequency was slightly higher in the TD group than the ASD group. In the sensitivity analysis, when the analysis was repeated on FOOOF parameters derived after exclusion of the six peripheral electrodes, these results remained unchanged.

**Table 4:** The estimated intraclass correlation coefficients (ICCs) and their 95% bootstrap confidence intervals for the six FOOOF parameters for each diagnostic group.

| FOOOF Parameter | TD | ASD |
| --- | --- | --- |
| Offset | 0.484 (0.004, 0.775) | 0.525 (0.167, 0.806) |
| Slope | 0.284 (0, 0.674) | 0.699 (0.527, 0.815) |
| Number of peaks | 0.081 (0, 0.571) | 0.226 (0.003, 0.609) |
| Largest alpha peak center frequency | 0.700 (0.437, 0.862) | 0.619 (0.342, 0.852) |
| Largest alpha peak amplitude | 0.862 (0.729, 0.939) | 0.828 (0.664, 0.926) |
| Largest alpha peak bandwidth | 0.424 (0.028, 0.696) | 0.340 (0.034, 0.727) |

### Nonparametric Analysis of the Relative Power Spectral Density via Functional Data Analysis

The estimated functional ICC for the scalp-wide relative spectral density was excellent in both groups, though higher in the TD group than the ASD group (TD 0.858 95% CI [0.748, 0.926]; ASD 0.807 95% CI [0.650, 0.914]). The estimated functional ICC for each of the 18 electrodes and their 95% bootstrap CIs are shown by diagnostic group in Figure 3. While the average electrode-specific ICC in the TD group is approximately equal to that of the ASD group, there is greater variation in the functional ICC among electrodes in the TD group (both higher and lower values of the functional ICC) compared to the ASD group. In the sensitivity analysis, the estimated scalp-wide functional ICC for both diagnostic groups was slightly higher when the six peripheral electrodes are excluded (TD 0.874 95% CI [0.741, 0.931]; ASD 0.815 95% CI [0.712, 0.913]), though the magnitude of difference between the two diagnostic groups was unchanged.

**Figure 3:**
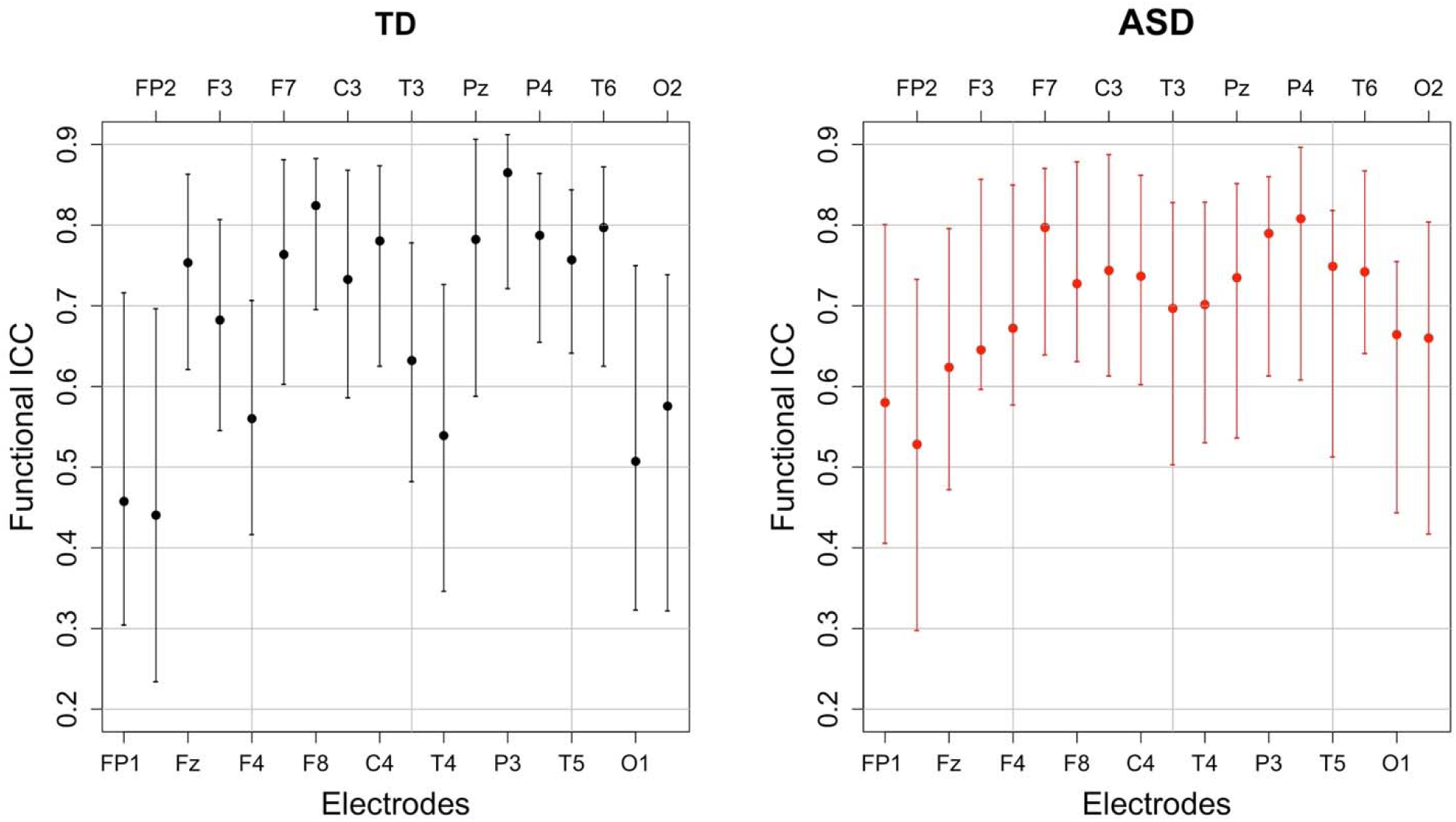
The estimated electrode-specific functional intraclass correlations and their 95% bootstrap confidence intervals by diagnostic group.

The functional ANOVA model captures individual deviations from the mean scalp-wide relative spectral density over the two days by partitioning the total variance into participant- and day-level variation. Participant-level variation captures the variation among participants whereas day-level variation captures the variation within a subject across days. Within each level of variation, ordered curves known as eigenfunctions identify which portions of the frequency domain account for the most variation by placing more magnitude at these locations. The two estimated leading participant- and day-level eigenfunctions for both diagnostic groups are shown in Figure 4. We restrict our discussion to the first two participant-level eigenfunctions, since combined they explain at least 60% of the total variation in both groups. We include the first two day-level eigenfunctions for completeness. The first participant-level eigenfunction for both groups displays that most variation in the data is explained by the variation in the amplitude of the alpha peak (with maximal variation at approximately 9 Hz), explaining similar total variation for the TD group (48% total variance explained) and the ASD group (43% total variance explained). While the first subject-level eigenfunction highlights variation in the amplitude of the largest peak, the second subject-level eigenfunction highlights the variation in the frequency (location) of the largest peak, where TD subjects show the largest variation in the low and high alpha band (24% total variance explained) and ASD subjects show it in high alpha and beta relative power (18% variance explained). These findings are consistent with the locations of the largest peak summarized in Table 3 across days for the two groups. The fact that most of the variation is explained by the subject-level eigenfunctions (compared to day-level eigenfunctions) supports our interpretation that most of the variation in the data is variation across subjects and there is less variability within a subject across days. In addition, participants maintain stable alpha peaks across experimental days, both in terms of peak frequency and amplitude, consistent with the high ICCs reported in Table 4 for alpha peak amplitude and frequency in the two groups in the FOOOF analysis.

**Figure 4:**
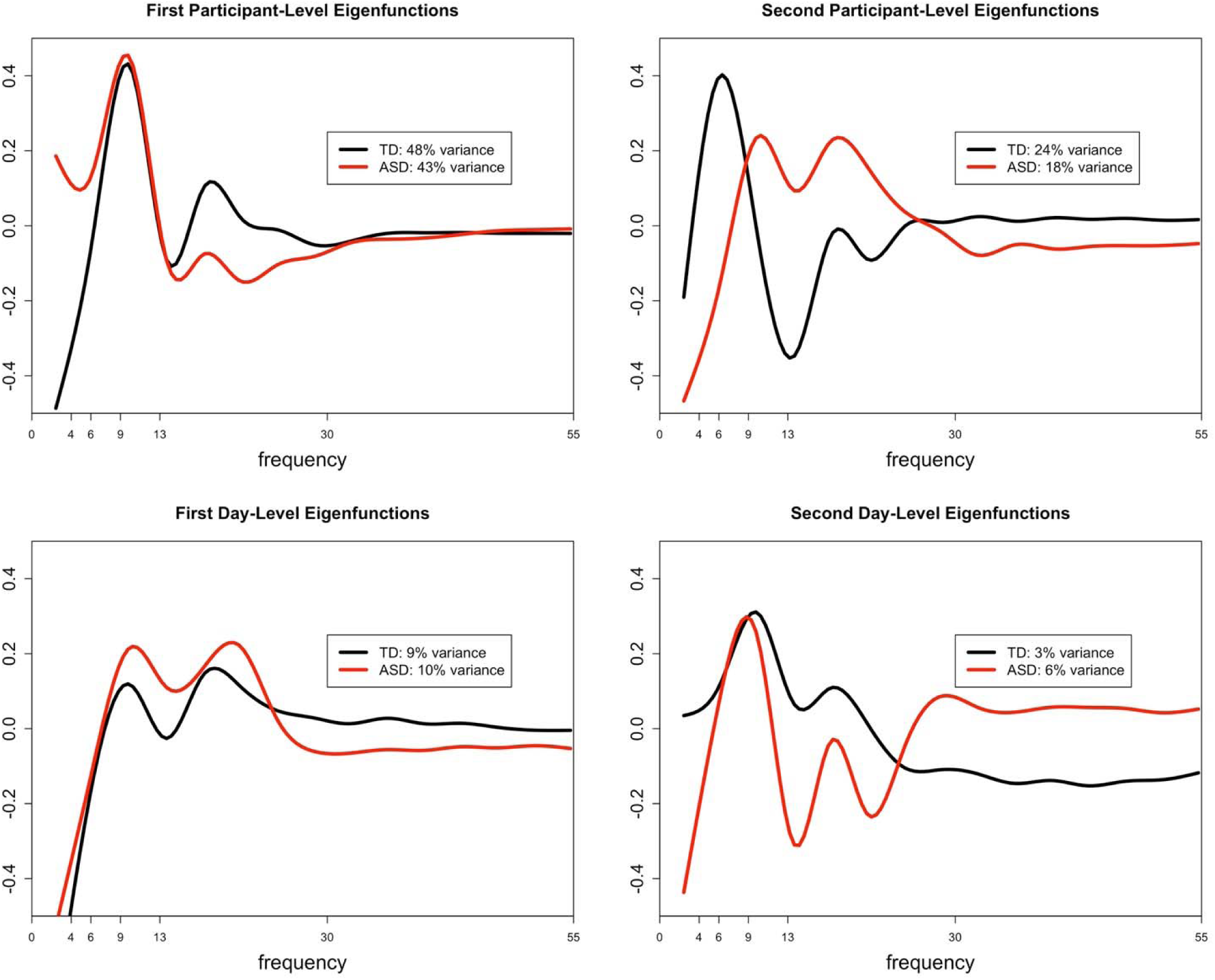
The estimated first and second leading eigenfunctions for the participant-level variation (top row) and day-level variation (bottom row) for each diagnostic group. The total variation explained by each component is included in the legend.

## Discussion

In this manuscript, we examine the test-retest reliability of the EEG power spectral density in children with ASD and TD. EEG power-based measures are currently being evaluated and employed as biomarkers in a variety of neurodevelopmental and psychiatric disorders, and analytical validation (including understanding the test-retest reliability of these measures) is an important early step in the biomarker development process.^35^

Overall, our findings demonstrate excellent test-retest reliability for scalp-wide EEG profiles. This high test-retest reliability reflects the overall stability of the EEG power spectrum over relatively short time windows (a few days). For the development of diagnostic biomarkers, this reliability is crucial – we would not expect the fundamental biology of the brain to change over several days, and therefore biomarkers indexing brain function for diagnostic purposes should not change significantly over this time period.

On the other hand, there are scenarios in which we would not expect (or want) aspects of the EEG power spectrum to remain stable. For example, while markers of phenotypic traits may remain stable, markers of state may vary over short time periods. For example, changes in emotional state during testing, and attention to the stimuli, may lead to changes in EEG power that reflect true physiologic changes in brain function over even short time windows. Identifying the parameters of the EEG PSD that predominantly reflect trait, and separately those that predominantly reflect state, will allow us to harness the wealth of information available from EEG recordings to develop a range of biomarker types. This concept will be crucial for future studies as well. For example, monitoring biomarkers will ideally remain relatively stable when treatment is not given, but show significant change in response to targeted medical treatments.

The high test-retest reliability for EEG profiles is present in both TD and ASD groups, though reliability was higher overall in the TD group (ICC 0.858) than the ASD group (ICC 0.807). This is consistent with prior findings suggesting more variable neural activity in ASD compared to TD^8^ and may suggest that reliability, in addition to providing important information for biomarker development, may in and of itself represent a potential biomarker. It is also possible that the lower mean IQ in the ASD group (or, perhaps less likely, the higher mean age of the ASD group) contributed to this difference Notably, higher neural variability may reflect (or provoke) more variable emotional states during testing and more variable attention to the stimuli. Such factors are often found to be clinically more variable among children with ASD.

Because the EEG PSD captures a range of parameters, it is important to consider specifically which of those parameters have high short-term test-retest reliability (and thus offer potential for diagnostic biomarker development), versus those with low short-term test-retest reliability (potentially reflecting state, attention or perhaps noise). Our findings suggest that within the PSD, a relatively small set of parameters are largely responsible for capturing the fingerprint-like quality of each individual’s EEG. FOOOF-based parameterization suggests that the alpha peak is particularly useful for individualizing the power spectrum. Within the alpha peak, amplitude offers particular promise in this regard, although the center frequency of the alpha peak also provides strong reliability within individuals. Here, it is particularly notable that the frequency of alpha is often considered to be an individual trait (changing only gradually with age and other factors but otherwise remaining relatively stable in most cases), whereas alpha amplitude varies more with state. For example, the posterior dominant rhythm tends to arise when the eyes are closed and is suppressed with eye opening; similarly, mu rhythms over the motor cortex are suppressed by imagining or engaging in motor tasks. However, our findings suggest that in the context of the environment in which EEGs were collected in the ABC-CT (watching silent, screen-saver type videos), alpha amplitude remains quite stable – even more so, in fact, than alpha frequency.

For the slope of the power spectrum as measured by FOOOF, ICC was good in the ASD group but poor in the TD group. This suggests that slope (at least as measured by FOOOF with the parameters used here) is unstable across sessions in the TD group. One possible explanation for this is that the TD group may be more sensitive to session effects (e.g., due to habituation, adaptation, or learning) than the ASD group, and this is being reflected in the slope. It is also possible that the older mean age or lower mean IQ of the ASD group, rather than TD or ASD status per se, contributed to this difference. An alternative explanation, supported by visual review of figure 2, is that there is very little inter-individual variability in the PSD slope among the TD group; therefore, intra-individual reliability (across days) cannot be much higher than inter-individual reliability (across participants) in the TD group, because inter-individual reliability is high to begin with. In the ASD group, which may be more heterogeneous given the wide variety of genetic and other underlying factors that lead to ASD, the inter-individual variability in slope is higher. In this case, similarly strong intra-individual reliability in the TD and ASD groups would lead to a higher ICC in the ASD group, because of the higher inter-individual variability in this group.

Importantly, the eigenfunctions which best characterized PSD shape exhibited the most variance at relatively low frequencies (4-13hz), corresponding to overall offsets of the PSD and in the theta to alpha range of the EEG, aligning with the parametric findings from FOOOF and highlighting the import of this frequency range for characterizing stable inter-individual differences in brain activity. This finding, combined with the tendency for variance to be explained by activity at slightly higher frequencies in the ASD group (alpha-beta) than TD participants (predominantly alpha), may help to explain the higher estimated ICC for offset and slope in the ASD group compared to TD. Because the slope and offset terms in FOOOF are fit in the semilog-power space, these parameters are sensitive to power dynamics at higher frequencies, which are often of lower magnitude.

For the nonparametric analyses of relative power, reliability in both groups improves with removal of peripheral electrodes. Notably, because peripheral electrodes are closer than central electrodes to many non-brain-based sources of detected activity (e.g., muscle and eye movements), they are often more susceptible to artifact than more central electrodes. This suggests (perhaps reassuringly) that brain-based findings, more so than artifact-based findings, remain stable across EEG sessions within an individual. On the other hand, for the parametric analyses of absolute power, removal of peripheral electrodes does not improve reliability. This may be because the majority of parameters identified by FOOOF are not significantly affected by artifact in peripheral electrodes, raising the possibility that FOOOF is less susceptible to artifact contamination than nonparametric analyses; this may be further studied in future work.

Nonparametric analyses otherwise reveal complementary results to the parametric analyses. Parametric analyses reveal excellent ICC for the amplitude of the largest alpha peak and good ICC for the frequency of the largest alpha peak. This is true in both the ASD and TD groups, though the ICC in the TD group is slightly higher than that in the ASD group for both of these parameters. Similarly, nonparametric analyses highlight alpha amplitude as capturing the majority of variance for the participant-level spectral densities, followed by alpha frequency. This is again true in both the ASD and TD groups, though slightly more variance is captured by the first two eigenfunctions in the TD as compared to the ASD group. Parametric functions also demonstrate that the dominant peak tended to stay within the alpha band for the TD group, but tended to stay more broadly in the range of both the alpha and beta bands for the ASD group. Similarly, nonparametric functions demonstrate that the TD participants show the largest variation in the alpha band, whereas ASD participants show variation in alpha but also extending into beta.

Nonparametric functional data analysis and FOOOF thus provide convergent and complementary approaches to characterizing the PSD. Nonparametric functional data analysis characterizes PSD shape accurately and with a small number of principle functions yielding high levels of reliability. However, it relies on “learning” these functions based on the current data set and thus yields different principle functions based on the input data, as we see here between our diagnostic groups. Additionally, the resulting functions need careful interpretation to ground their relationship with brain activity. Conversely, FOOOF estimates require more parameters to characterize the PSD. However, fitting these parameters does not depend on the presence of other members of the data set, (although the algorithm fitting settings can indirectly force information sharing among power spectra). Also, the interpretation of FOOOF parameters is more direct. FOOOF explicitly attempts to separate biophysically meaningful model parameters such as slope, offset, and oscillatory peaks.

It is important to note the specific questions that the present study is designed to answer. First, the two testing days for each individual took place within approximately a week. While this suggests promise for biomarker development in trials where EEG-based findings are expected to change over very short periods of time, many pharmacological interventions aim to change neural activity over the longer term (weeks, months, or longer). Examining test-retest stability of the EEG power spectrum over these longer periods is part of ongoing analyses for the ABC-CT main study, which will include 6 week and 6 month follow-up recordings. Additionally, here we report only test-retest reliability for a single set of EEG measures, all based on the power spectrum. EEG is a rich source of information beyond that which can be captured in the power spectrum, in both the time domain and the frequency domain. As future studies suggest additional EEG-based measurements that may offer promise for biomarker developments, the test-retest reliability of the measurements will need to be explicitly evaluated.

Developing biomarkers for ASD and other neurodevelopmental disorders remains a high priority in the field, given the potential benefits biomarkers offer for clinical trials, diagnostics, and monitoring.^4^ While future studies will continue to assess which measurements (in EEG and otherwise) offer the most promise as potential biomarkers of various types, our findings of high short-term test-retest reliability of the EEG power spectral density are a crucial step towards ensuring that potential biomarkers meet necessary criteria for validation.

## Author Contributions

- ARL, AJN, AS, SJW, FS, CS, MM, RAB, KC, GD, SF, SJ, CAN, JCM, and DS made substantial contributions to the conception or design of the ABC-CT.
- ARL, AJN, AS, and DS contributed to the analysis of the data described in this manuscript.
- ARL, AJN, AS, SJW, and DS contributed to the drafting this manuscript.
- ARL, AJN, AS, SJW, FS, CS, MM, RAB, KC, GD, SF, SJ, CAN, JCM, and DS provided critical revisions related to the important intellectual content.
- All named authors read and provided approval for publication of the content.

## Acknowledgments

A special thanks to all of the families and participants who join with us in this effort. In addition, we thank our external advisor board, NIH scientific partners, and the FNIH Biomarkers Consortium.

Additional important contributions were provided by members of the ABC-CT consortium including Heather Borland and Megha Santhosh, who were responsible for EEG acquisition including EEG experimental and pipeline programming, site training and initiation, and quality control.

